# The genetic structure within a single tree is determined by the behavior of the stem cells in the meristem

**DOI:** 10.1101/2022.11.07.515471

**Authors:** Yoh Iwasa, Sou Tomimoto, Akiko Satake

## Abstract

Genomic sequencing revealed that somatic mutations cause a genetic differentiation of the cells in a single tree. In this study, we consider a mathematical model for stem cell proliferation in the shoot apical meristem (abbrev. SAM), which results in genetic diversification between the cells differing in the distance along the shoot and the angle around a shoot axis. The assumptions are as follows. Stem cells in the SAM normally undergo asymmetric cell division and produce successor stem cells and differentiated cells. The differentiated cells proliferate and contribute to shoot elongation.

Occasionally, a stem cell is replaced by a copy of an adjacent stem cell. We discuss the “coalescent length” between cells indicating their genetic difference with respect to neutral mutations. A mathematical analysis revealed the following. The genetic diversity of cells sampled at the same position along the shoot increases with the distance from the bottom of the shoot. Stem cells hold a larger variation if they are replaced only by the nearest neighbors than if they are replaced by any cells. The coalescent length between two cells increases not only with the difference in the position along the shoot but also in the angle around the shoot axis. The dynamics of stem cells at the SAM determine the genetic pattern of the entire shoot.

## 1. Introduction

In a long-lived terrestrial plant, such as a tree, somatic mutations produce genetic and epigenetic differences between cells within a single individual. Mutations may be caused by errors in genomic duplication or from the recovery from a physical and chemical damage to the genome. Recent genomic sequencing techniques, such as next-generation sequencing, provide all the genome information of the cells at different portions of a tree, revealing the pattern of genetic diversity pattern within an individual plant (Schmidt-Siegert 2017; Plomion et al. 2018; Hanlon et al. 2019; Orr et a. 2020; Hofmeister et al. 2020; Yu et al. 2020). The accumulation of mutations occurs as the shoot elongates. We expect that two positions of a single shoot (trunk or branch) could exhibit genetic differences in a magnitude correlated with their physical distance along the shoot.

The genetic diversity within an individual tree has been discussed by theoretical population geneticists (Antolin and Strobeck 1985; Klekowski and Kazarinova-Fukshansky 1984a; Tomimoto and Satake 2022). Many of these works focus on the effect of natural selection at the cellular level, the genetic loads, mutation meltdowns, and the enhanced potential of adaptation in a changing environment (Klekowski and Kazarinova-Fukshansky 1984b; Otto and Orive 1995; Pineda-Krch and Fagerström 1999; Folse and Roughgarden 2012). Others have discussed the potential effects of the layer structure in the meristem (Klekowski et al. 1985; Pineda-Krch and Lehtilä 2002). Although knowing the outcomes of selection operating at the cellular level is biologically important, measuring cellular fitness in a quantitative manner is still difficult. In contrast, the recent rapid advancements in the genomic sequencing techniques has made it rather easy for us to obtain the spatial distribution of genetic variation within an entire tree, although obtaining the genome from a single cell is still difficult (Reusch et al. 2021).

Some studies have noted the correspondence between the molecular phylogeny of cells within a tree and the physical shape of a tree (Orr et al. 2020; Zahradníková et al. 2020). To clarify the logic, Sou Tomimoto and Akiko Satake (unpublished) introduced a distinction between “path after forking” and “path before forking.” Suppose that two cells, A and B, on different branches are sampled, as illustrated in Fig. 1A. The shortest path connecting A and B along the trunk and branches is called the path after forking. Because the genetic difference between the two cells is an outcome of mutation accumulation in shoot elongation, it is expected to increase with the length of the path after forking. In Fig. 1A, the shoot elongation leading to A, and that leading to B, originated at the fork marked as F. Because the shoot apical meristem (abbrev. SAM) includes multiple stem cells, the ancestor stem cells of A and B at F can be different. For an extreme case, if A and B were descendants of different stem cells at the start of the entire shoot, the common ancestor of A and B should be in the bottom of the shoot, rather than at the forking F. Then, the genetic difference between A and B should be explained by the mutation accumulation in the path before forking and the path after forking combined (see Fig. 1A). The importance of mutation accumulation in the path before forking depends on the stem cell dynamics within the SAM (as explained below).

**Fig. 1.**
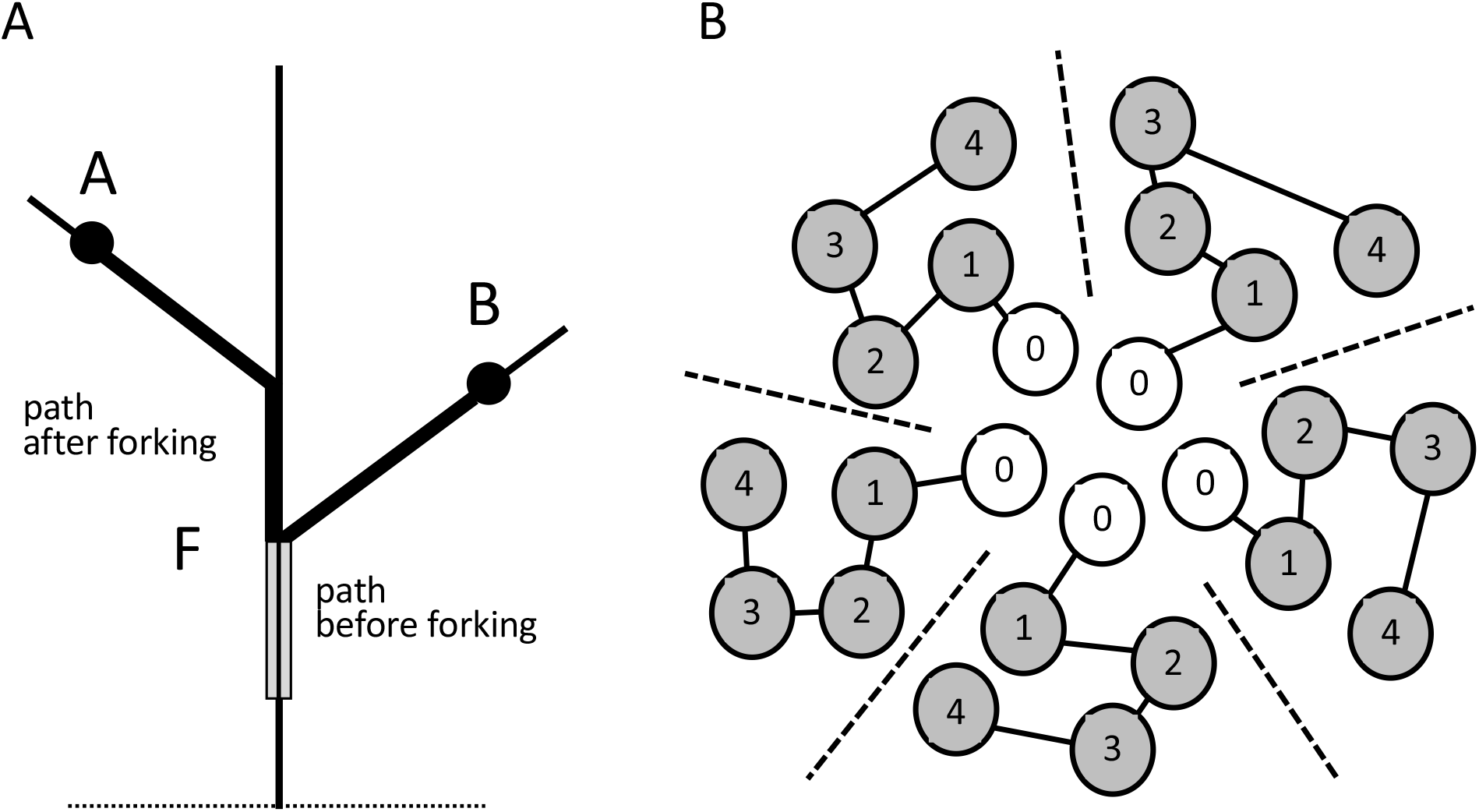
Schemes for the model. (A) Paths after forking and before forking of a shoot elongation. Cells A and B are sampled from two different branches. The location of the fork is indicated by F. The genetic differentiation between the two cells is caused by the mutations occurring in the shoot elongation of both “after forking” and “before forking”. See text for explanations. The concepts were introduced by S. Tomimoto and A. Satake (manuscript). (B) Scheme of shoot apical meristem. The white circles indicate stem cells and shaded circles indicate differentiated cells. The broken lines separate five columns. Stem cells undergo asymmetric cell division and produce a successor stem cell and a differentiated cell. Differentiated cells increase in number within each column.

To quantify the genetic structure of an entire individual tree, we can focus on the similarity between cells sampled from the different portions of a tree. This is equivalent to coalescent theory, that is useful in understanding the geographical genetic structure formed by different processes, such as migration, habitat heterogeneity, episodic selection events, population bottleneck, and the reproductive isolation from the data of genetic similarity of different subpopulations or different taxonomic units (Kingman 1982, Hudson 1983).

In this study, we attempt to develop a coalescent theory for an individual tree. Specifically, we focus on the coalescent process between different cells constituting a single shoot. We aim to clarify the processes generating the before-forking portion of the path, although we do not handle branch formation in this paper. All the cells constituting a shoot originate from the stem cells at the SAM. The spatial size of the SAM is on the order of 100 *μ*m (or 0.1 mm). The cell division, replacement, and mobility in this small tissue determines the genetic structure of the entire tree, which may have a size on the order of 10 meters.

If the stem cells at the shoot apical meristem (SAM) undergo asymmetric cell division (Fig. 2A), they leave their own successors without failure, and they accumulate different mutations from the start of shoot elongation. As the shoot grows, stem cells at the SAM become genetically different. However, a stem cell may occasionally fail to leave its successor by performing a symmetric division leaving two differentiated cells (Fig. 2B). If this happens, another stem cell duplicates to recover the number of stem cells at the SAM, which was inferred from the empirical studies of chimeric plants (Stewart and Dermen 1970; Ruth et al. 1985). In mathematical analyses of stem cell dynamics, the stem cell was assumed to be replaced by a cell randomly chosen from the remaining stem cells (e.g., Klekowski and Kazarinova-Fukshansky, 1984a). This assumption is plausible for the haematopoietic stem cells of mammals (e.g. Marciniak-Czochra et al. 2009).

**Fig. 2.**
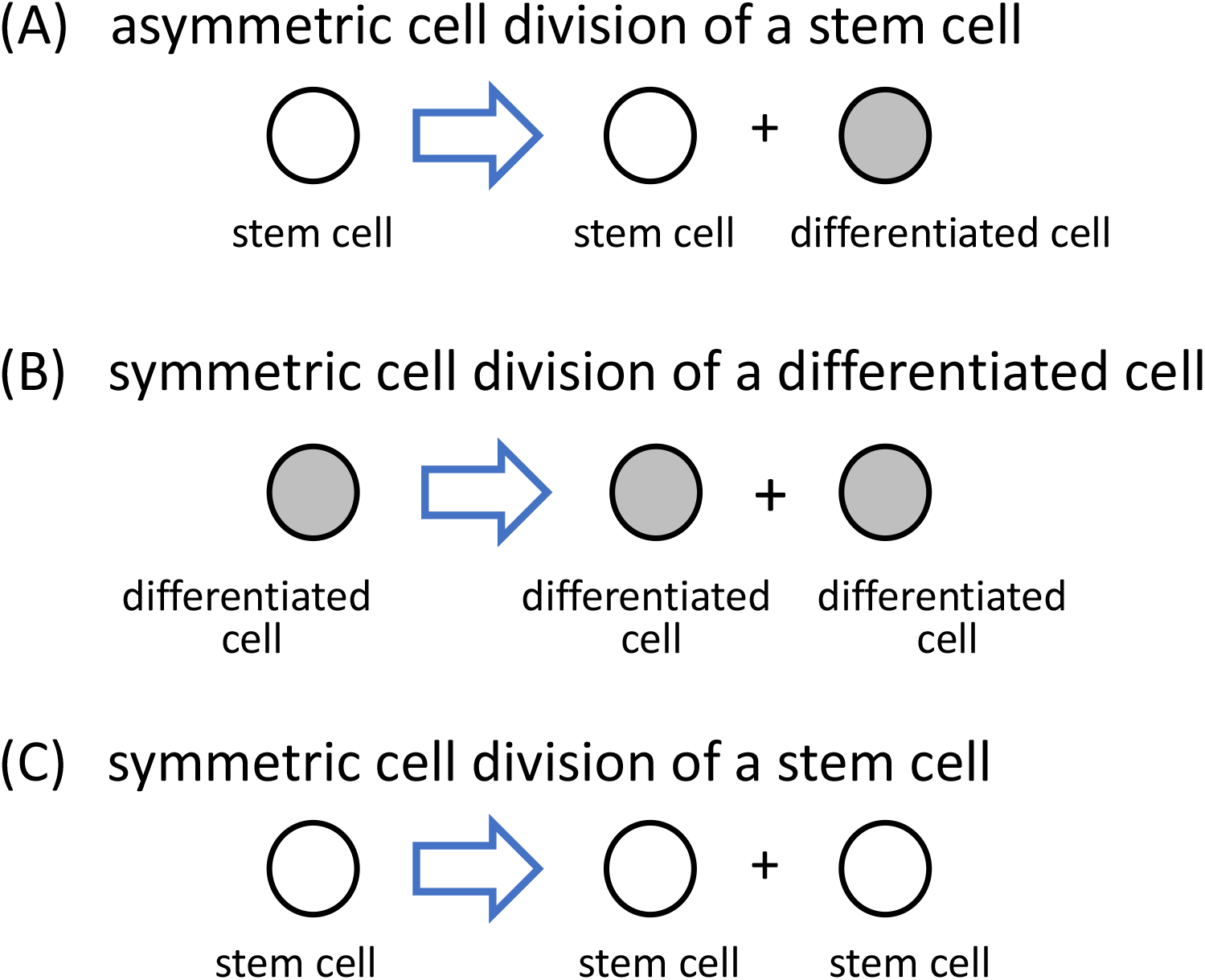
Three modes of cell division. (A) Asymmetric cell division of a stem cell, producing a stem cell and a differentiated cell. (B) Duplication of a differentiated cell, producing two differentiated cells. (C) Duplication of a stem cell. This occurs only when another stem cell fails to produce its own successor.

However, plant cells have cell walls that make it difficult for them to exchange locations. Here, we consider an alternative assumption: the vacancy of stem cells should be filled by the duplication of one of the neighboring stem cells, located either to the left or right of the lost stem cell (see Fig. 1B). This assumption results in the genetic structure around the axis of the shoot: two stem cells become genetically more similar if they are similar in angle than if they are located on the opposite side of the shoot. Stem cells in the SAM develop a genetic structure related to the angle around the shoot axis in a manner similar to the circular stepping-stone model (Maruyama 1969, 1970a, 1970b, 1971), which was developed to handle the “isolation by distance” concept in population genetics theory (Wright 1943; Kimura and Weiss 1964). With this spatial structure, the stem cells at the SAM maintain a larger total genetic variation than the case when stem cells are replaced irrespective of their locations.

To discuss how different stem cell dynamics produce different genetic patterns of somatic mutations, we analyzed the “coalescent length” between cells, which indicates their genetic difference in regards to neutral mutations. A mathematical analysis revealed the following. The genetic diversity of cells sampled at the same position along the shoot increases with the distance from the bottom of the shoot. Stem cells hold a larger variation if they are replaced only by the nearest neighbors than if they are replaced by any cells. The coalescent length between two cells increases not only with the difference in their position along the shoot but also in the angle around the shoot axis. The dynamics of the stem cells at the SAM determine the genetic pattern of the whole shoot.

## 2. Model

Fig. 1B illustrates a scheme of a shoot apical meristem (SAM) viewed from the above. The five open circles indicate stem cells. They normally engage in asymmetric cell division: a randomly chosen stem cell divides and produces a successor stem cell and a differentiated cell (shaded circle). The differentiated cell is placed in a sector close to the parental cell. The broken lines in Fig. 1B indicate separations between sectors. Fig. 2A explains the asymmetric cell division, which makes the number of stem cells remain constant. A differentiated cell that is newly formed by the asymmetric cell division of a stem cell subsequently repeats symmetric cell divisions for a finite number of times, as illustrated in Fig. 2B. This, together with the increase in cell size, results in shoot elongation. Consequently, a sector of the SAM in Fig. 1B forms a “column” of the shoot. The differentiated cells in different columns are not mixed.

Fig. 3 illustrates the correspondence between asymmetric stem cell divisions at different times and the descendant differentiated cells existing at different positions of the shoot. In Fig. 3A, the differentiated cells numbered 8, 9, 10, and 11 were produced from the asymmetric cell division at time *t* = 1; the cells numbered 4, 5, 6, and 7 were from asymmetric cell division at time *t* = 2; the cells numbered 2 and 3 were from asymmetric cell division at time *t* = 3; and a differentiated cell numbered 1 was produced from asymmetric cell division at time *t* = 4. This shows that the physical position of the differentiated cells along a shoot axis corresponds to the time of the asymmetric cell division of the stem cell from which they originated. Differentiated cells in the lower (upper) position of a shoot can be regarded as a “fossil record” of the stem cells from earlier (later) days. We can determine the accumulation of mutations occurring in the stem cell line by comparing cells at different positions of a shoot. In Fig. 3B, the SAM (white ellipsoid on the tip) moves upwards, contributing to shoot elongation. The lower (upper) portions of the shoot were formed from the stem cells in an earlier (later) time.

**Fig. 3.**
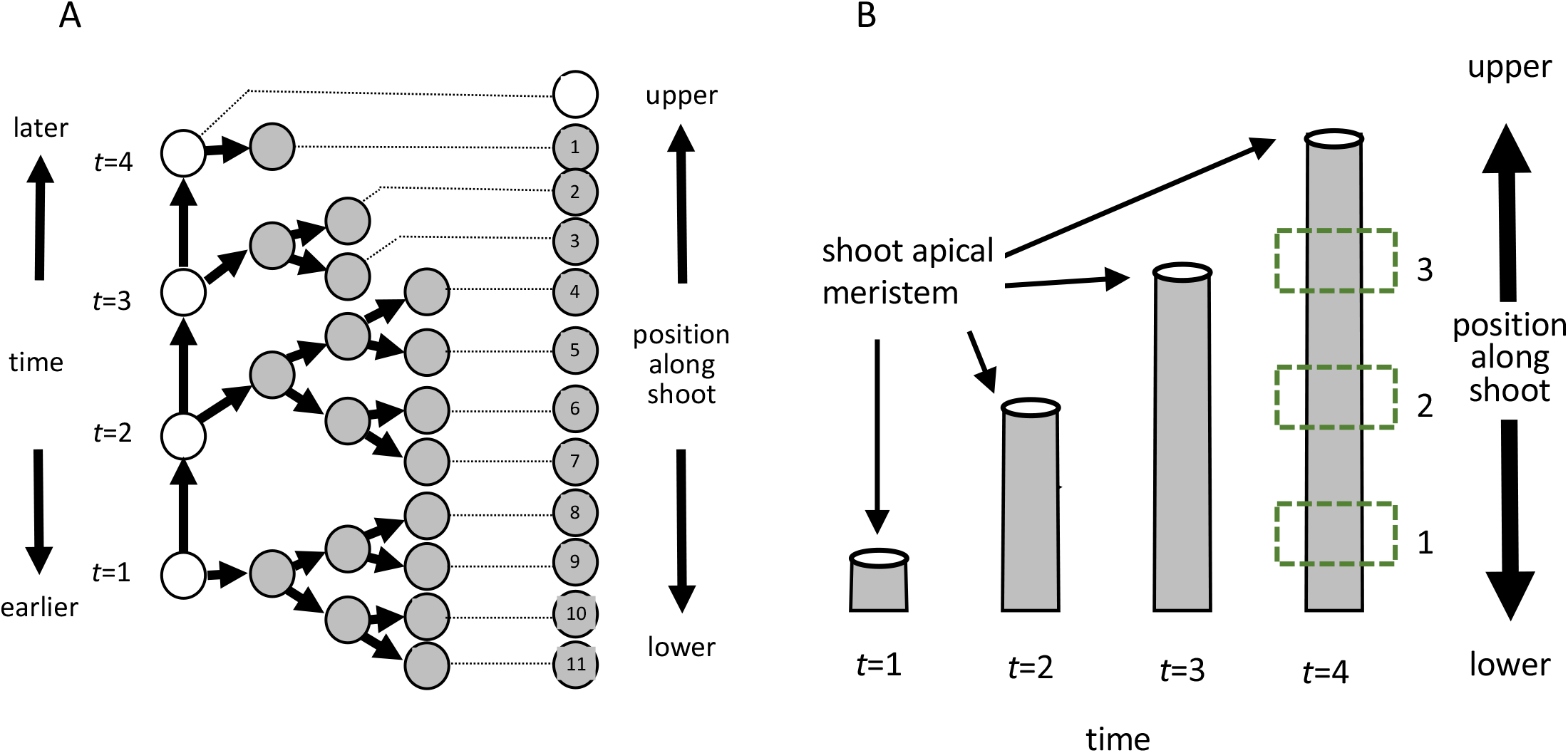
Correspondence between the time at which a stem cell undergoes an asymmetric cell division and the physical location of the descendant differentiated cells along the shoot. (A) The left part indicates four asymmetric cell divisions of a stem cell (*t* = 1,2, 3,4). The right part indicates the location of the differentiated cells along the shoot. Cells in the low position (8, 9, 10, 11) are descendants of an asymmetric cell division at an earlier time (*t* = 1). In contrast, cells in the upper position (e.g., 2, 3) are descendants of an asymmetric cell division at a later time (*t* = 3). (B) The stems are at different time points. White ellipsoids on the tip of the shoot indicate the shoot apical meristem at which the stem cells engage in asymmetric cell division producing differentiated cells. As time goes by, the shoot grows and the location of the meristem moves upwards. The shaded portion of the shoot indicates differentiated cells. The rightmost shoot is the one at time *t* = 4. The three dotted squares numbered 1, 2, and 3 indicate the portions of the shoot formed by the descendants of the asymmetric cell division at time *t* = 1,2, and 3, respectively.

If stem cells engage in asymmetric cell division only, the number of stem cells remains the same as at the start of shoot starts. Here, we assume that stem cells at the meristem have the same genome when the shoot started. As the shoot grows, the lineages of the stem cells in the SAM become differentiated from each other as different stem cells accumulate novel mutations independently. Consequently, the genetic diversity among stem cells in the SAM keeps increasing.

### 2.1 Failure of asymmetric cell division and the replacement of stem cells

However, occasionally a stem cell may fail to leave a successor daughter stem cell. Then, to recover the stem cell number at the SAM, one of the nearest neighbors duplicates, and fills the site of the stem cell, as illustrated in Fig. 2(C). This “stem-cell replacement” event results in the loss of a stem cell lineage and the reduction of genetic diversity in the stem cell population. The balance between the accumulation of novel mutations and the loss of genetic diversity by stem cell replacement makes the average genetic diversity among the stem cells in the SAM converge to a stationary level.

All stem cells exist in the center of the meristem (Fig. 1A). Cells in the plant tissue are not as mobile as those in animal tissues owing to their cell wall. For each stem cell, the nearest stem cells are those either on the left or on the right. It is plausible to consider that stem cell replacement occurs between the nearest neighbor cells arranged in a circular manner. For illustration in Fig. 1A, the five sections of cells correspond to the five angles around the shoot axis, which leads to five columns of the shoot.

In general, there are *n* columns arranged in a circular manner (*k* = 1,2,*n*). The left and right nearest neighbors of column *k* are columns *k* – 1 and *k* + 1, respectively. Owing to the circular boundary condition, to the right of column *n* is column 1. We adopt modular arithmetic with a modulo *n* calculation with respect to the symbol for the column number. We assume that *n* columns are distributed evenly over 2*π* (360 degrees), with column *k* facing towards the angle of 2*πk/n*. For the sake of simplicity, we call *k* the “angle around the shoot axis” in the calculations below. We assume that the stem cell placed in column *k* can be replaced by a copy of the stem cell in the left column (column *k* – 1) or in the right column (column *k* + 1) respectively, at rate of *c* per unit time. The replacement of a stem cell line by a neighboring stem cell line causes mixing between the neighboring columns, leading to a genetic similarity of neighboring columns. Consequently, the stem cells develop a circular genetic structure; cells with closer angles around the shoot axis (close in circular column number *k*) are more similar than those with distant angles. The model has aspects common to the circular stepping-stone model for geographically structured populations, which was developed to handle the “isolation by distance” concept (Maruyama 1969, 1970a, 1970b, 1971).

### 2.2 Coalescent length

In molecular phylogeny, the genetic differentiation between two genomes is measured in terms of the coalescent time, which is twice the time from their common ancestor (Kingman 1982; Hudson 1983). Because the physical length along the shoot is easier to measure than the time difference between two events of stem cell division in the meristem, we here consider the “coalescent length” to quantify the potential for holding the genetic difference between the different parts of a shoot.

A differentiated cell sampled in an empirical study and its ancestral stem cell at the time of asymmetric cell division differ because of the mutations in the proliferation of the differentiated cells (Fig. 3A). In the following calculation, we focus on the distance between the stem cells from which two sampled cells originated. To compare the predictions of the model with the data, we need to consider the additional length caused by the mutations occurring between the sampled differentiated cells and the stem cells from which they are derived.

As illustrated in Fig. 3, the distance between two positions along the shoot corresponds to the difference in the time at which the cells were produced by asymmetric division of a stem cell. If the speed of shoot growth is constant, the distance between the two sampled positions along the shoot can be converted to the difference in the time when the two portions were produced from stem cells in the meristem. A stem cell divides and produces a differentiated cell at rate *r* (asymmetric cell division). Each newly produced differentiated cell ends up with a number of cells by symmetric cell division for a finite number of times (Fig. 2B), which occupy a portion of the shoot with physical length *g*. We call *g* as the “shoot growth rate per asymmetric cell division.” Time interval Δ*t* makes a stem cell produce differentiated cells that occupy the shoot length *rg*Δ*t*. Hence, the shoot length of Δ*z* corresponds to the time interval of length Δ*z/rg*.

The genetic differences between cells sampled at the same position can be measured in terms of the “coalescent length between cells,” defined in the following manner. As illustrated in Fig. 4, we always have a single ancestor of the focal cell at all the time points before the sampled time (i.e., all the position below the sampled point). An occasional failure to leave a successor stem cell results in stem cell replacement, which occurs at a rate of *c* per unit time. If this occurs, the site for occupied by the stem cell is filled by a copy of either the left stem cell or the right stem cell with an equal chance. When we trace the ancestral lineage of a cell, this appears as a “column shift” (Fig. 4). For a randomly chosen stem cell, in a short time interval Δt, the lineage experiences the column shift with the left stem cell and with the right stem cell with probability 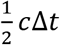. Hence, *q* = *c/rg* is the factor converting the unit length of a shoot to the probability for a column exchange event for a stem cell. We call *q* as ‘‘the column shift rate per unit length of shoot.”

**Fig. 4.**
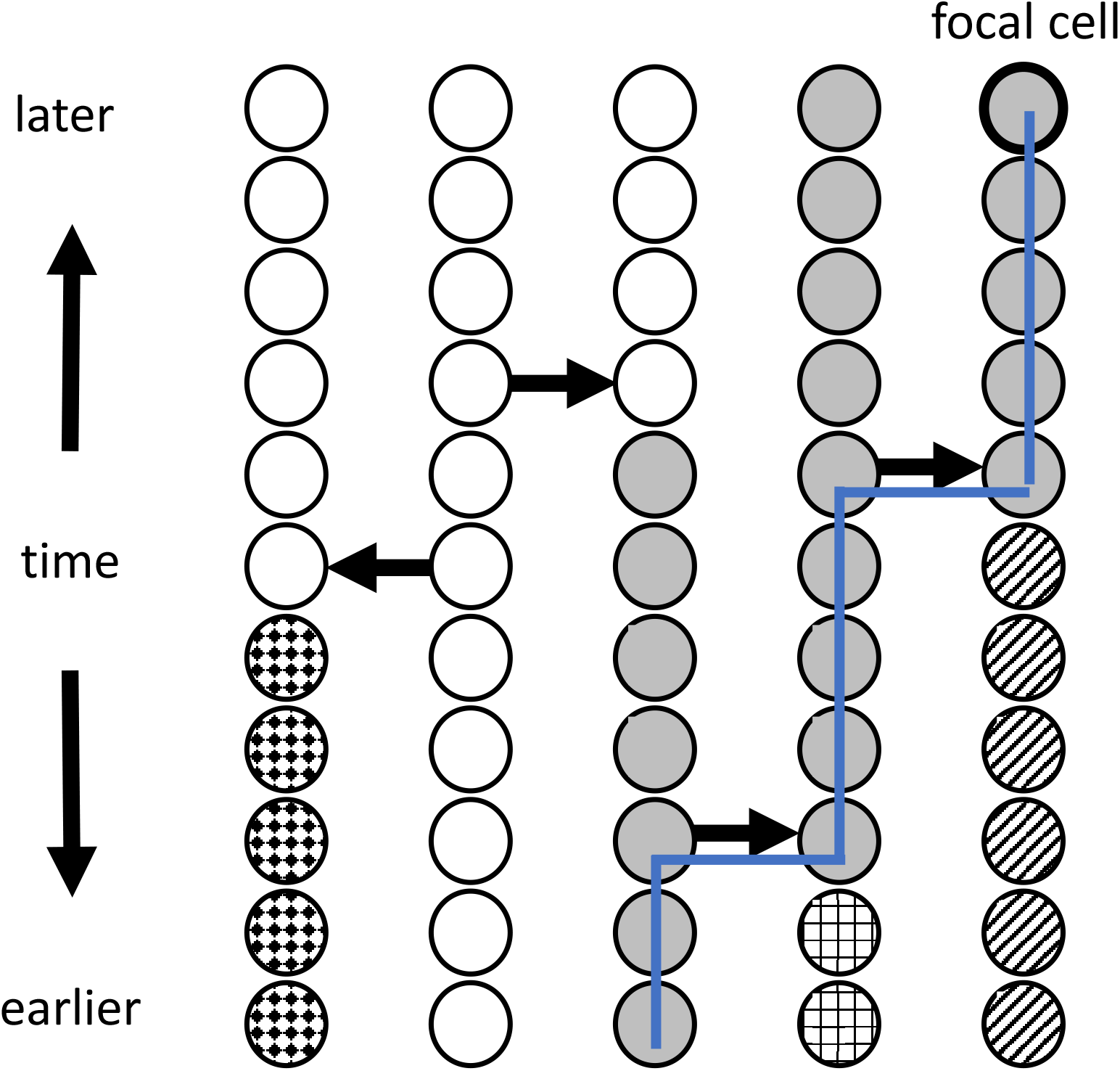
Ancestral lineage of stem cells. The five columns of circles indicate the stem cells in different columns. The vertical location indicates the time, with the lower and upper positions indicating the earlier and later times, respectively. The horizontal arrows indicate the stem cell replacement events. When a stem cell fails to produce a successor stem cell, a copy of one of the neighboring stem cells fills the site. The pattern of circles indicates the lineage of the stem cells. If we choose a focal cell at the final time, we can trace the ancestral cell lineage. One example of the ancestral cell lineage of the focal cell (the rightmost cell at the final time) is shown by a blue line. There were the five different cells at the initial time (the lowest layer) but there were descendants of only two cells among the five existing in the final time (the top layer). This implies that the stem cell replacement reduces the genetic variation of the stem cells. On the other hand, cells receive novel mutations, which makes the cells different from each other genetically (not shown here).

Suppose that we trace the ancestral lineages of two different cells sampled at the same position of a shoot. Their ancestral lineages should move randomly between the neighbors among *n* stem cells (Fig. 5). Due to random movement, their ancestral cells may overlap with each other when they have a common ancestor. This implies that the coalescence of the two sampled cells occurs, and the doubled length of time between the time of coalescence and the time of sampling is the coalescent time (Kingman 1982; Hudson 1983). If we convert the time to the physical length along the shoot, the coalescent time becomes the coalescent length.

**Fig. 5.**
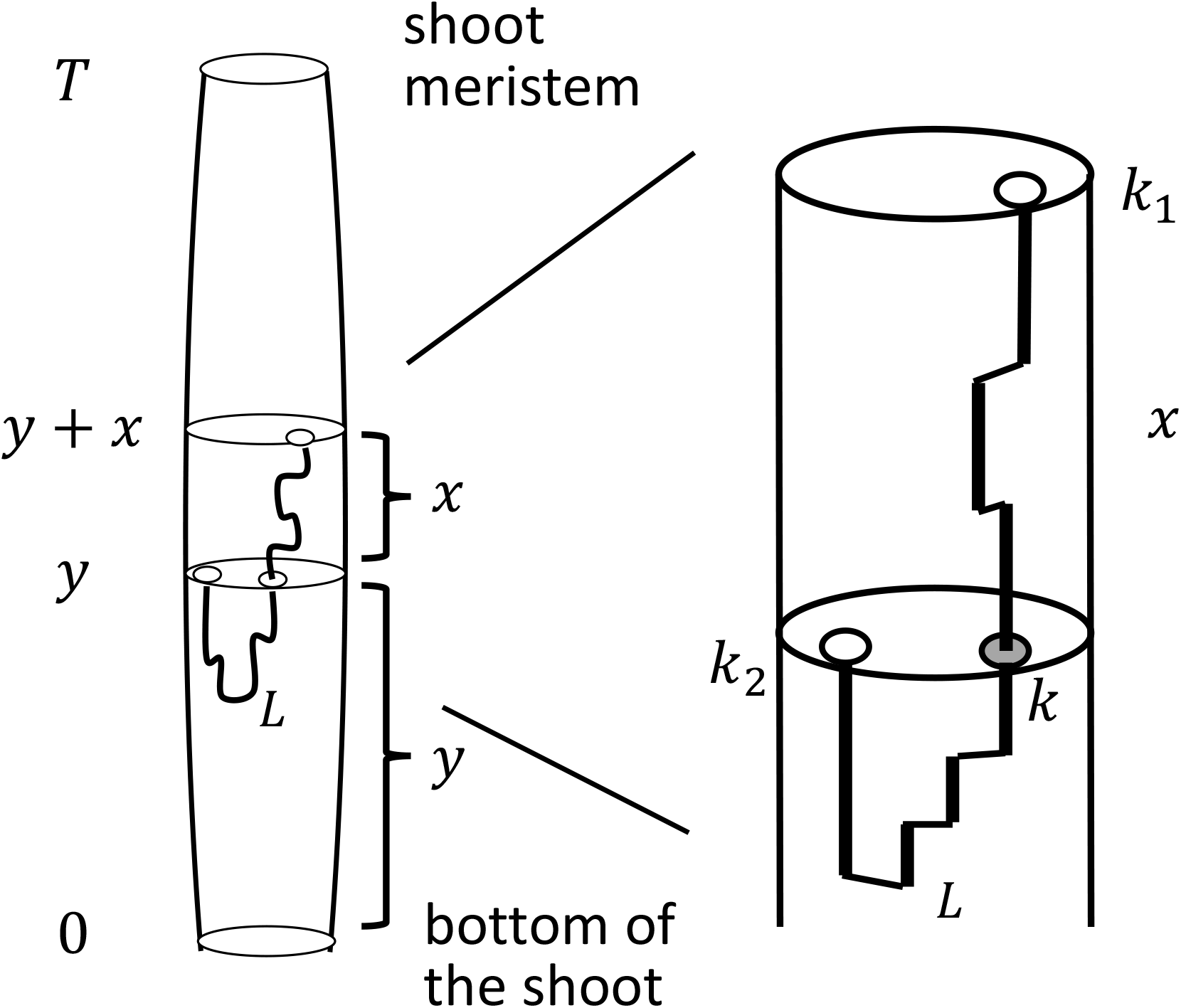
Scheme of coalescent length. A cell at position *y* + *x* and angle *k*_1_, and the second cell at a position *y* and angle *k*_2_ are sampled. The ancestral cell of the former exists at a position lower than *y* + *x*. It has angle *k* at the position *y*. The ancestral cell of the first sampled cell and the second cell may not be the same (*k*_2_ ≠ *k*). Then we need to calculate the expected coalescent length between them. It is *L*_*k*–*k*_2__(*y*).

We indicate the position along a shoot by *z*, which is defined as the distance from the bottom of the shoot. The differentiated cells with the positions of the smaller *z* (lower position) reflect the genetic state of the stem cells at an older time, and those with the positions of the larger z (upper position) reflect the state of the stem cells at a more recent time. Let *T* be the length of the whole shoot. *z* = *T* is the current position of the SAM where the stem cells are proliferating. Let *z* = *y* be the position at which we sampled two cells (Fig. 5). For a stem cell sampled at a time, at any time earlier than it, there existed a single cell that was the ancestor of the focal cell. Tracing the ancestral cell for a chosen cell is a symmetric random walk between columns. Note that the opposite does not hold. If we pick up a cell at one time, at any time later than it, there can be no descendant, or two or more descendants.

We consider a cell at position *z* in column *k* and consider the events experienced by its ancestral cell occurring between *z* – Δ*z* and *z*, where Δz is a short distance along the shoot. Note that we trace an ancestral cell lineage from the descendants to the ancestors. For each cell in a column, a column shift may take place within a short distance Δ*z* with probability *q*Δ*z*, where *q* = *c/rg* is the rate of stem-cell replacement per unit shoot length. If a column shift occurs in the focal column within interval (*z* – Δ*z, z*), the newly formed cell at *z* is a daughter of the cell either in column *k* – 1 (the left one) or in column *k* + 1 (the right one).

## 3. Coalescent length between two cells sampled at the same position

Cells sampled at the same position of a shoot can be genetically heterogeneous because they were formed from the genetically heterogeneous stem cells at the SAM. We assume that the stem cells were genetically homogeneous at the start of the shoot growth.

In Fig. 4, five columns are illustrated as five vertical lines. Because of the periodic boundary condition, the leftmost column is the next to the rightmost column.

The top of the vertical lines is the cell group at the focal position. If we trace the lines downwards, we have positions lower along the shoot. The horizontal arrows represent stem cell replacement events, each of which indicates that the cell in the column at the time was a daughter cell of a neighboring stem cell (either right or left). After a stem cell replacement event, the genetic difference between the two cell lineages becomes zero.

Let *L_k_*(*y*) be the mean coalescent length between two cells sampled at the same position. This depends on *y*, the distance from the bottom of the shoot, and the angle difference between the two sampled cells. We consider the events occurring in a short period corresponding to a short physical interval between *y* – Δ*y* and *y*. We have the following recursive formula:

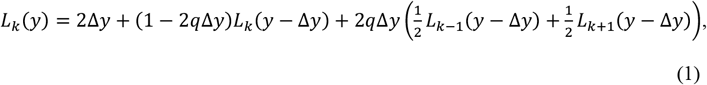

The left-hand side is the mean coalescent length of two cells in columns with their angle difference *k*. The first term on the right-hand side of Eq. (1) indicates the increase in the coalescent length 2Δ*y*, which is the doubled length of the interval Δ*y*, because there are two branches. The second term indicates the probability of having no event in this interval (1 – 2*q*Δ*y*) multiplied by the mean coalescent time *L_k_*(*y* – Δ*y*). The third term indicates the probability that one of the two ancestral cells experiences a replacement 2*q*Δ*y* multiplied by the coalescent length in such situations. When a stem cell replacement by the left or right stem cell occurs, the angular difference either decreases (column difference *k* – 1) or increases (column difference *k* + 1) with equal probability.

By rearranging terms, dividing both sides by Δ*y*, and calculating the limit Δ*y* → 0, Eq. (1) becomes

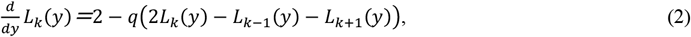

where *k* = 1,2,.., *n*; *y* > 0. Because sampling from the same column would result in a zero distance, we have *L*_0_(*y*) = *L_n_*(*y*) = 0 for *y* > 0. This implies that we have an absorbing boundary condition. Because there is no genetic diversity at the bottom of the shoot, the initial condition is *L_k_*(0) = 0 for *k* = 1,2,..,*n* – 1. By solving these differential equations numerically, we obtain the coalescent length *L_k_*(*y*).

### 3.1 Mathematical results for several simple cases

We can obtain an explicit mathematical solution for *L_k_*(*y*) in simple cases, as explained in Appendix A.

As indicated in the boundary condition for Eq. (2), the mean coalescent length is short near the bottom of the shoot. It increases with *y* and converges to the value given by the following formula:

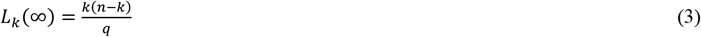

where *q* = *c/rg*. See Appendix A for the derivation. Eq. (3) indicates that the coalescent length depends on *k*, the difference in column number between the sampled cells, or the angular difference, as illustrated in Figs. 6A and 6B. It is small when the two cells are similar in angle (small *k*) and it is large when they are different. The maximum value of the coalescent length, given in Eq. (3), is *n*^2^/4*q*, when the two sampled cells are from the opposite side of the axis (*k* ≈ *n*/2). The mean value of the coalescent length when all *k*’s have the same probability is (*n*^2^ – 1)/6*q*. Hence the genetic diversity among the cells at the same position of the shoot can be large when the number of stem cells at the shoot apical meristem is large (large *n*) and when each stem cell undergoes asymmetric cell division without failure (small *q*).

**Fig. 6.**
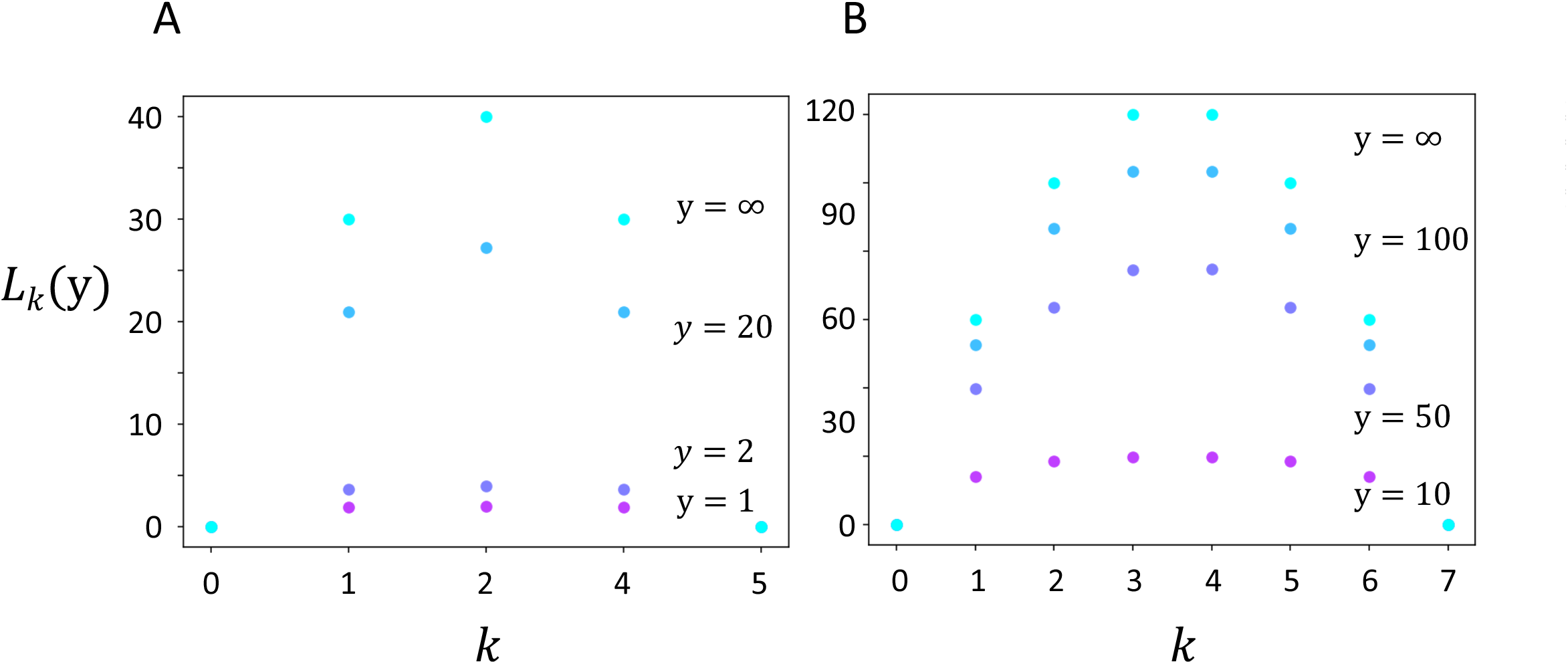
Coalescent length. The vertical axis represents the expected coalescent length *L_k_*(*y*), and the horizontal axis represents the column number (or angle) *k*. Different curves are for different values of *y*, the length from the bottom of the shoot. (A) *n* = 4. (B) *n* = 7. The other parameter was *q* = 0.1.

Fig. 6A illustrates the *L_k_*(*y*). It is small when *y* is small, and it increases monotonically with *y*. This implies that the cell diversity should increase as we move from the lower to the upper positions along the shoot. The value at a sufficiently far distance from the bottom, given by Eq. (3), depends on *k*, the angle difference between two sampled cells. It is small when the two samples are on close in angle and is at the maximum when the two sampled cells are on the opposite side of the shoot.

When no replacement of stem cells occurs (*q* = 0), two cells in different columns have a common ancestor only at the beginning of shoot formation. Hence, the coalescent length is the maximum value *L_k_*(*y*) = 2*y* (if *k* ≠ 0).

For the general value of *y* > 0, we can calculate the solution *L_k_*(*y*) explicitly using Laplace transforms, as explained in Appendix A. As an illustrating example, we consider the case with *n* = 4. We rewrite Eq. (2) using matrix notation as follows:

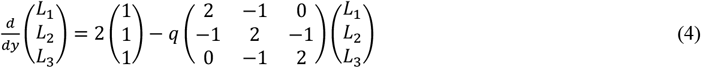

From the boundary condition for Eq. (2), we have *L*_0_(*y*) = *L*_4_(*y*) = 0. Note that the matrix is of a 3 × 3 form (instead of 4 × 4). After some calculations, we have

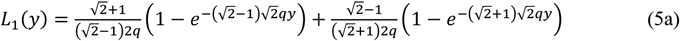

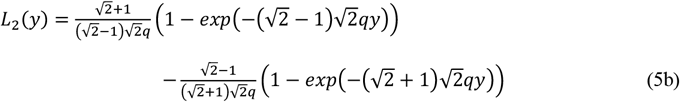

From symmetry, *L*_3_(*x*) = *L*_1_(*x*) holds. We can confirm these results by a numerical calculation of differential equations.

In Appendix A, we explain that, for any *n, L_k_*(*y*) can be obtained using the inverse Laplace transforms. We also show explicit solutions for *n* = 2, *n* = 3 and *n* = 4. However, for the solution for *n* = 5 or a larger *n*, the solution is messy. We obtain the solution by a numerical analysis of the differential equations given in Eq. (2).

## 4. Coalescent length between two cells sampled at different positions

Suppose we sample two cells at positions *y* and *y* + *x* as illustrated in Fig. 5. Note that *y* + *x* and *y* are the upper and lower positions of a shoot, respectively. We traced from the upper cell to its ancestral cell, which exists in any position lower than it. They are separated by *x*, and their angles around the shoot axis were *k*_1_ and *k*_2_, respectively. We consider the ancestor of the cell sampled at position *y* + *x*, and we denote that the angle of the ancestor cell at position *y* is in column *k* (Fig. 7). We denote the probability for the ancestral cell to be in column *k* by *Q*_*k*–*k*_1__(*x*). It depends on *x*, the distance between the two cells along the shoot. If *x* is very small, the probability distribution *Q*_*k*–*k*_1__(*x*) has a sharp peak at *k* = *k*_1_. If *x* is larger, the distribution *Q*_*k*–*k*_1__(*x*) becomes flattened, and in the limit *x* → ∞, it becomes a uniform distribution over *n* sites, where all the columns have an equal chance of having the ancestral cell.

**Fig. 7.**
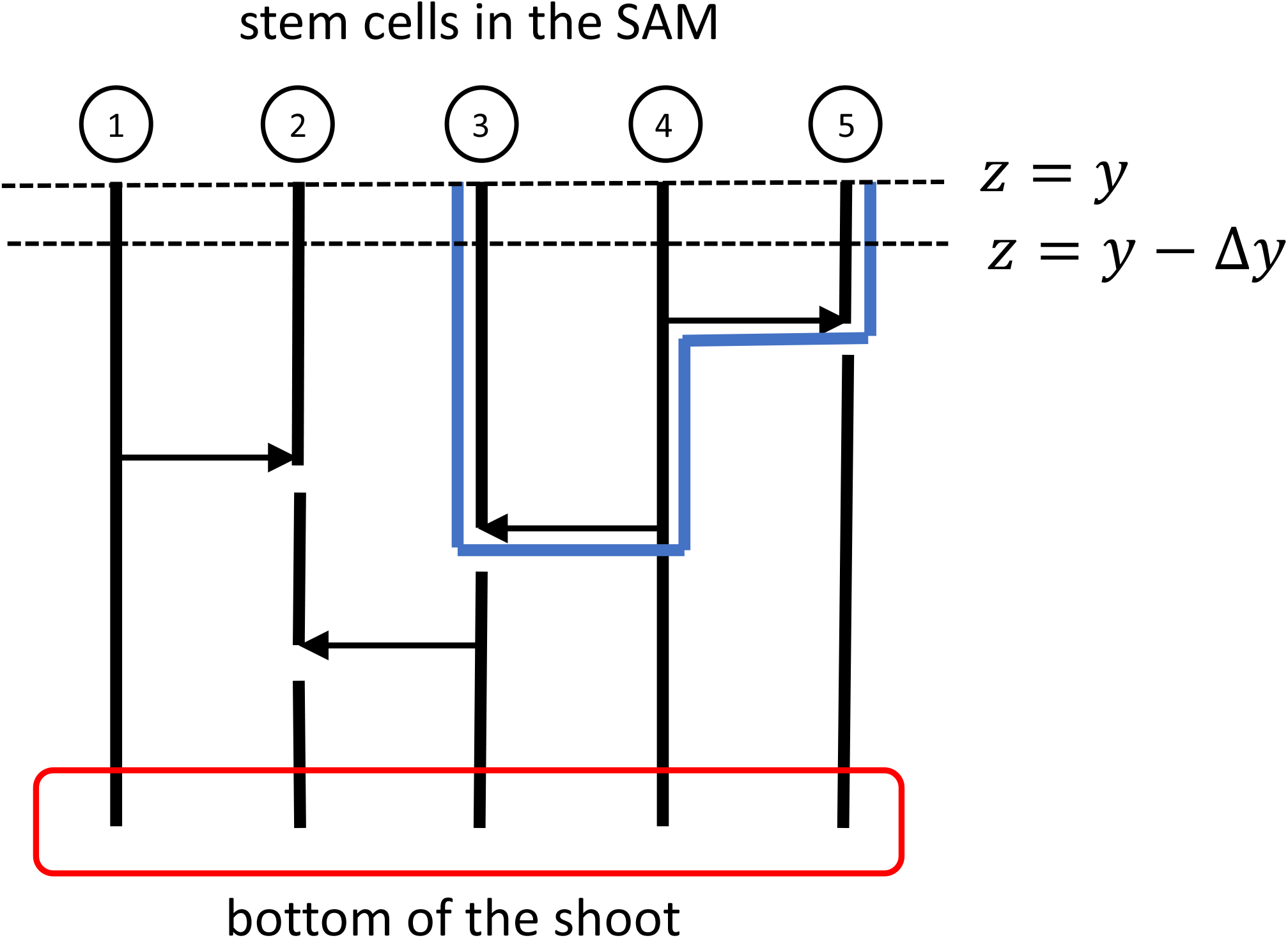
Coalescent time. The five vertical lines indicate fine stem cells, and the vertical axis is for the time (the lower and upper locations are for the earlier and later time points, respectively). The horizontal arrows indicate the stem cell replacement (see caption to Fig. 4). The ancestral lineages of cell numbers 3 and 5 are indicated by blue lines. They have a common ancestral cell among the middle of the vertical line. Coalescence occurs at this point, and the coalescent time is defined as the length of the line connected between two sampled cells (3 and 5 in this illustration), which is the doubled time since the coalescence. In the current paper, we evaluate the coalescent length, which is the value converted from the coalescent time.

The mean coalescent length between the ancestor cell *k* and the second sampled cell *k*_2_, both being at position *z* = *y*, is *L*_*k*–*k*_2__(*y*), which was discussed in the last section. The mean coalescent length between two cells sampled at different positions is the sum of two terms:

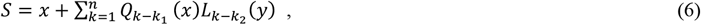

where the first term on the right-hand side is the distance between the two sampled positions along the branch. The second term is the mean coalescent length indicating the genetic variation among the cells at the same position. It is calculated as the average of *L*_*k*–*k*_2__(*y*) with probability *Q*_*k*–*k*_1__(*x*), (*k* = 1,2,.., *n* – 1).

### 4.1 Probability distribution of the ancestor cell

Here, we consider how the distribution of ancestor cells changes with *x* and the distance along the shoot. Suppose we sample a stem cell at one position along the shoot and consider its ancestral cell at the position lower than the sampled cell by the distance *x*. Let *Q_k_*(*x*) be the probability for the ancestor cell to be in column *k*. We have the initial distribution: *Q*_0_(0) = *Q_n_*(0) = 1; and *Q_k_*(0) = 0 for *k* ≠ 0, because the focal cell existed in column 0, which is column *n*. We have the following formula:

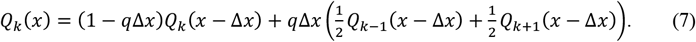

The term on the left-hand side indicates the probability for the ancestor cell to be in column *k* at position below the focal descendant cell by a distance of *x*. We decompose this probability according to the events that occur between *x* – Δ*x* and *x*. The first term on the right-hand side is the probability of the absence of a stem-cell replacement (1 – *q*Δ*x*) multiplied by the same distribution at position *x* – Δ*x*. *q* = *c/rg* is the stem-cell replacement rate per unit shoot length. The second term corresponds to the case in which the cell was a daughter of one of the nearest neighboring stem cells with probability *q*Δ*x*. The brackets include two terms corresponding to the columns of the parental cell being either column *k* – 1 or column *k* + 1, with equal probability.

By rearranging terms and dividing both sides by Δ*x*, and calculating the limit Δ*x* → 0, we have the following differential equation:

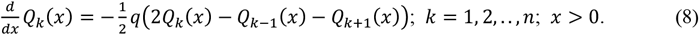

A system of *n* differential equations given in Eq. (8) with the initial condition can be solved explicitly using matrix calculations, as explained in Appendix B. We can obtain the probability distribution of the ancestor cell among columns, *Q_k_*(*x*), for all *k* and *x*, as follows:

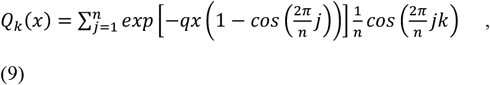

where *q* = *c/rg*. When *x* is small, the distribution has a sharp peak at *k* = 0. The height of the peak declines with *x*, and the probability distribution becomes more evenly distributed. Fig. 8 illustrates how *Q_k_*(*x*) changes with *x* and *k*. Note that *Q_k_*(*0*) satisfies the initial condition, and in the limit *x* → ∞, 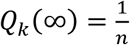 holds for all *k*. For an intermediate *y*, we can confirm Eq. (9) by a numerical calculation of the differential equations, given in Eq. (8).

**Fig. 8.**
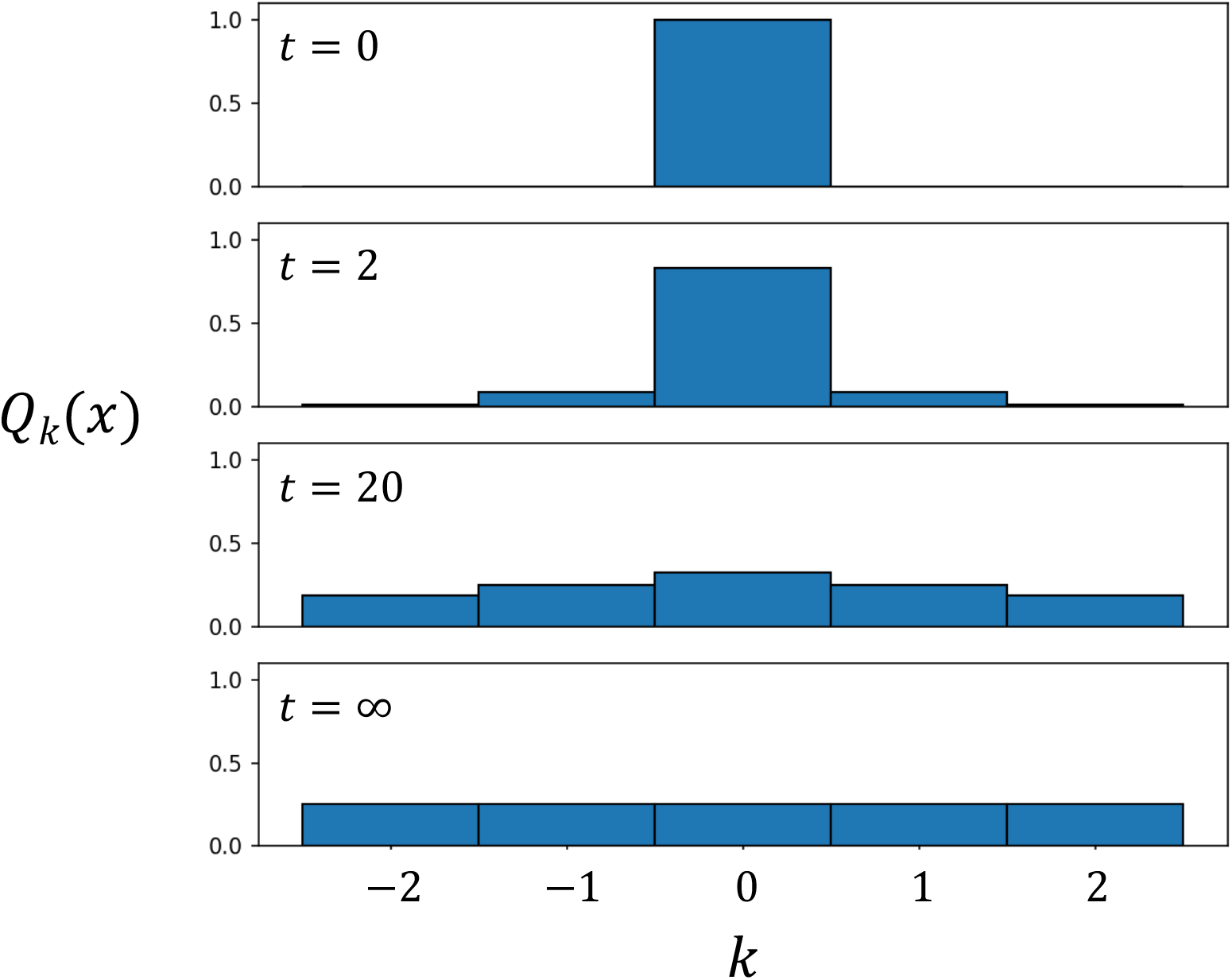
The probability distribution of the location of the ancestral cell, *Q_k_*(*x*). The horizontal axis is the column number *k*, which is expressed in terms of modulo *n*. The results for several different values of the distance *x* are shown. When *x* = 0, the probability is concentrated to *k* = 0, which is the same as *k* = *n*. As *x* increases the peak becomes lower and the distribution becomes more widespread. In the limit *x* → ∞, all the columns have equal probability. The graph is calculated from Eq. (9). The parameters are: *n* = 4, *q* = 0.1.

### 4.2 Mean coalescent length between two cells sampled at different positions

Suppose that one cell is sampled at position *y* + *x*, and the second cell is sampled at position *y* with their angles *k*_1_ and *k*_2_, respectively. Owing to the homogeneity of the process, the mean coalescent length between the two cells depends only on the physical distance of the two positions, and the difference in the two angles (*x* and *k*_1_ – *k*_2_).

If the sampled position is sufficiently far from the bottom of the shoot (*y* →∞). Eq. (6) becomes

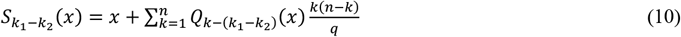

using Eq. (3). The second term on the right-hand side is the magnitude of the coalescent length caused by the genetic diversity among the cells at the same position.

When *qx* ≪ 1, *Q*_*k*–(*k*_1_–*k*_2_)_(*x*) concentrates on *k* ≈ *k*_1_ – *k*_2_, then the second term is close to (*k*_1_ – *k*_2_)(*n* – (*k*_1_ – *k*_2_))/*q*. When *k*_1_ = *k*_2_, it is zero. When |*k*_1_ – *k*_2_| ≈ *n*/2, it is approximately *n*^2^/4*q*. Hence, the relative position of the two sampled cells is important. The genetic difference is larger when they are sampled from the opposite side of a shoot than when they are from the same side of the shoot.

When *qx* ≫ 1, *Q*_*k*–(*k*_1_–*k*_2_)_(*x*) ≈ 1/*n*, which is independent of the relative position of the two sampled cells (*k*_1_ – *k*_2_). In addition, the effect of the first term is larger than that of the second term in Eq. (10). We can simply regard that the main determinant of the coalescent length is the first term *x*, the physical distance along the shoot between the two sampled cells.

## 5. Comparison with the case without spatial structure

In the current study, because stem cell replacement occurs only by the nearest neighbors, stem cells develop a circular structure of genetic similarity. In theoretical models for the intraindividual genetic diversity of trees, a stem cell can be replaced by a daughter of a randomly chosen stem cell in the meristem, which is the same as the continuous-time version of the Moran model in theoretical population genetics (Nowak 2006), although other models assume that all the stem cells are replaced simultaneously following the Wright-Fisher model (Crow and Kimura 1970). The authors of these studies concluded that the stem cell diversity was rather small.

We analyzed the Moran model in Appendix C. In the Moran model, in the limit *y* → ∞, the coalescent length between two different stem cells is *L* ≈ (*n* – 1)/*q*. The average coalescent length between two randomly chosen stem cells is *L* ≈ (*n* – 1)^2^/*nq*, because the coalescent length is 0 if the two sampled cells are identical. This result should be compared with the result given in Eq. (3), which depends on the angular difference between the two sampled cells. It has the maximum max_k_*L_k_*(∞) ≈ *n*^2^/4*q* and the average over *k* is 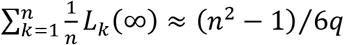. The average value of the coalescent length is greater than the corresponding value in the Moran process. Hence, we can conclude that the assumption of a stem cell replacement occurring between the nearest neighbors in the circular cell arrangement can enhance the genetic diversity maintained in the stem cell population at the meristem. The ratio of the average coalescent length between the two is *n*(*n* + 1)/6(*n* – 1), which increases with the number of stem cells (the ratio is 1.111 for *n* = 4; 1.4 for *n* = 6; 2.037 for *n* = 10). This effect is equivalent to the one found in the enhanced ability of maintaining a larger genetic variation in a geographically structured population more than in a perfectly mixed population, which can be represented in terms of a greater effective population size in a geographically structured population (Maruyama 1970a, 1971).

The effect of a differentiation between angles can be evaluated by considering the exchange of locations between stem cells. Suppose that stem cells in the shoot apical meristem can be mobile. If so, the neighboring stem cells may exchange their locations. Let s be the rate of the cell exchange event per unit time. If the exchange of their locations occurs sufficiently fast, the angle structure should disappear. However, the exchange of stem cells does not produce a coalescence between the stem cells. In Appendix D, we can derive the coalescent length when both a stem-cell replacement by a neighbor at rate *c*, and the stem-cell exchange between the neighbors at rate *s*, occur. The probability distribution {*Q_k_*(*x*); *k* = 1,.., *n* – 1} follows the same formula if we replace *c* with *c* + *s*. The calculation is given by Eq. (8) if *q* is replaced by *q*′ = (*c* + *s*)/*rg*. When the mixing between stem cells is very fast, *s* is very large, the structure related to the angle around the shoot axis disappears quickly and 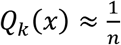 holds. In tracing ancestral cell lineages, column shifts occur frequently.

On the other hand, the calculation for *L_k_*(*y*) requires more care. In Appendix D, we can derive the following explicit formula for the coalescent length when the sampled position is sufficiently far from the shoot origin *y* → ∞.

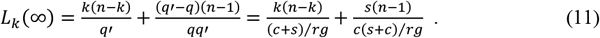

for *k* ≠ 0. This becomes Eq. (3) when the exchange is slow (*s* = 0). In contrast, when the stem cells are very mobile and the exchange rate is fast (*s* → ∞), Eq. (11) becomes 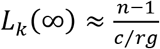. It is independent of angle *k* (for *k* ≠ 0), indicating that the genetic distance between the different stem cells becomes independent of the angle. It can be large if the stem cell replacement rate *c* is small. This result is the same as that for the Moran model because *q* = *c/rg*. The coalescent length when stem cells are very mobile (i.e., the Moran process) is smaller than in the angle-structured model studied in the current paper (*s* = 0).

## 6. Discussion

Terrestrial plants, such as long-lived trees, might develop a large genetic variance within an individual body because of somatic mutations. This has attracted research attention since the latter half of the last century (Whitham and Slobodchikoff 1981; Klakowski and Kazarinova-Fukshansky. 1984a, 1984b; Antolin and Strobeck, 1985; Sutherland and Watkins 1986; Otto and Orive 1995). In this paper, we analyzed a mathematical model by using the coalescent length and demonstrated that the genetic patterns developed by somatic mutations could be strongly affected by the behavior of stem cells at the shoot apical meristem (SAM). Formation of the axial meristem of a branch can be made by sampling a small number of stem cells of the main shoot meristem (Tomimoto and Satake 2022). If so, the genetic diversity of the stem cells at the SAM would contribute to the difference in the genetic compositions between the branches that are formed from the same shoot, in addition to the mutations occurring after the formation of the different branches.

In the models studied in the present work, there are two major processes that can generate a high genetic diversity between stem cells. First, stem cells in the shoot apical meristem undergo asymmetric cell divisions through which they leave their successor stem cells and differentiated cells. If the probability for stem cells to have successor stem cells is high, the stem cell lines are conserved from the beginning of the shoot and become genetically diverse as they accumulate different mutations independently. This should make the genetic diversity among the stem cells keep increasing with the distance from the bottom of the shoot.

Second, genetic diversity among the stem cells depends on how the stem cell replacement takes place when a stem cell fails to leave a successor stem cell. If the failed stem cell is replaced by the nearest neighbors, as we assumed in this paper, a circular similarity structure among stem cells is formed. The genetic difference between the stem cells should increase with the difference of stem cells in regards to the angle, and the total amount of genetic variation kept in the stem cell population is higher than in the case when the replacement occurs by randomly chosen stem cells, as assumed in other theoretical works.

We derived several predictions that can be tested by empirical studies.

1. Genetic diversity among cells at the same position of the shoot can be large if a stem cell replacement occurs very rarely (small *c*).
2. The genetic difference between cells may increase with the distance from the bottom of the shoot. The variation among differentiated cells at the upper position (large *y*) is larger than that at the lower position (small *y*).
3. Two cells with different angles (different *k*) around the shoot axis tend to be genetically more different than cells of similar angles (similar *k*).
4. Finally, stem cell dynamics at the shoot apical meristem determine the genetic structure of the entire shoot. The results depend on how accurately they can produce daughter stem cells, how often they are replaced by the other stem cells, which of the other stem cells replaces a cell that failed to succeed, and how mobile are the stem cells in the meristem.

In short, to understand the genetic structure of the entire shoot, we need to study the dynamics of the stem cells at the meristem carefully. The formula we presented in the present paper reveals the logical connections between the macroscopic pattern of the genetic differentiation of cells in an entire shoot and the microscopic cellular behavior within the SAM.

In comparing the predictions of the model in this paper and the observed genetic differentiation between the cells in different positions of an individual plant (such as a tree), we need to pay attention to the additional source of genetic variation. Differentiated cells experience cell divisions a finite number of times after their ancestral first differentiated cell was produced from a stem cell, as illustrated in Fig. 3A. If so, Eqs. (1) and (9) have an additional term that is independent of *x* or *n*.

In this study we focused on the genetic differentiation of cells along a single shoot but did not address branch formation, the latter being discussed by Tomimoto and Satake (2022). To model axillary bud formation for a new branch, they introduced the probability distribution of stem cells in the SAM of the trunk to contribute to the axillary meristem as well as a strong bottleneck. By incorporating these concepts, we may extend the mathematical analysis of coalescent length developed in the present paper to a situation where the sampled cells are on different branches rather than on a single shoot.

We discussed how the coalescent length depends on the distance between two sampled positions, the difference in the angle of two sampled cells, the distance from the bottom of the shoot, and the rate parameters describing the behavior of the stem cells at the SAM. Subtle differences in stem cell behavior at the SAM greatly affect the genetic structure of the entire shoot of a large tree. On the other hand, by observing the genetic structure of the cells in the shoot especially with respect to the longitudinal distance and angular difference, we may be able to obtain important information on the cellular behavior in the SAM. The correspondence between the genetic diversification at the whole-plant level and the cell mobility and replacement in the SAM is captured by mathematical formulas which should be useful in guiding research on the genomic structure and mosaicism of the whole plant in the coming years. Recently, sequencing technology has advanced very rapidly. We expect that the genomic sequence of a single cell will soon be available and will allow us to test all the predictions made in this study (Luo et al. 2020). We hope the current paper will stimulate efforts for more empirical studies in the near future.

## 7. Acknowledgements

This work was done in support of Kakenhi (Gants-in-aid for Scientific Research, JSPS): 17H06478, 20H00470, 21H04781 to A.S. We thank the following people for their very helpful comments: A. Hara, R. Hayashi, R. Imai, and E. Sasaki.

## Appendix A

We consider the following system of differential equations:

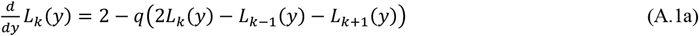

defined for *k* = 1,2,..,n – 1, and *y* > 0

The absorbing boundary condition is:

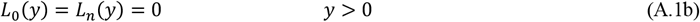

and the initial condition:

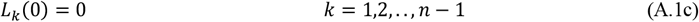

We note that we are handling a system of *n* – 1 differential equations. The boundary condition is absorbing at both *k* = 0 and *k* = *n*.

We consider the following square matrix of size (*n* – 1) × (*n* – 1):

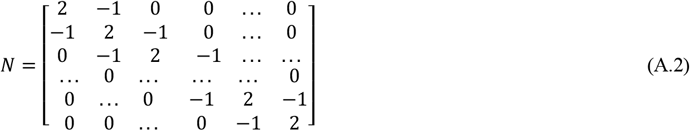

All the diagonal elements are 2, and the elements just below diagonal elements, and just above the diagonal elements are −1. All the other elements are 0. Using this matrix, Eq. (A.1) can be rewritten as

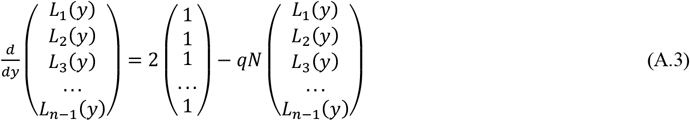

Note that the coefficient matrix *N* is different from *M* in Appendix A. The diagonal elements are 2, the elements just below the diagonal elements, and those just above the diagonal elements are −1, and all the other elements are 0. This is different from matrix *M* in Appendix A, in which both (1, *n*)- and (*n*, 1)-elements are −1. The eigenvalues and corresponding eigenvectors, or the inverse matrix, are obtained only for small *n*. Below, we show several cases for which we have some results.

### When y is very large

In the limit of *y* →∞, we have:

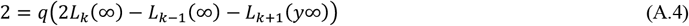

for *k* = 1,2,..,*n* – 1. The absorbing boundary condition is *L*_0_(∞) = *L_n_*(∞) = 0. We obtain the following function:

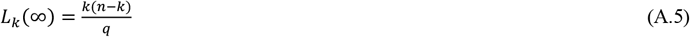

for *k* = 1,2,..,*n* – 1. We can confirm that Eq. (A.5) satisfies Eq. (A.4) and the boundary condition Eq. (A.1b).

### When y is very small

In the opposite limit of a very small *y*, we have

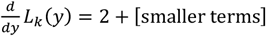

which can be solved as:

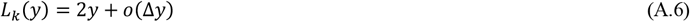

Between these two limiting cases (A.5) and (A.6), we expect that *L_k_*(*y*) monotonically increases with *y* according to numerical analyses.

For a few simple cases, we can solve the differential equations (A.1) explicitly.

### when n = 2

Because *L*_0_(*y*) = *L*_2_(*y*) = 0, the only equation we need to solve is:

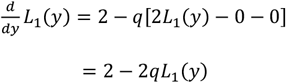

We can solve this with the initial condition *L*_1_(0) = 0, as follows:

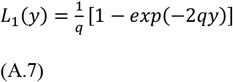

### When n = 3

When *n* = 3, according to the symmetry, we have *L*_1_(*y*) = *L*_2_(*y*). Noting that *L*_3_(*y*) = *L*_0_(*y*) = 0,

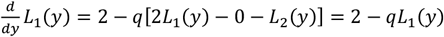

From this, we can solve the equation as follows:

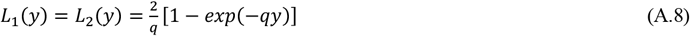

### when n = 4

When *n* = 4 or larger, the solution includes multiple time constants. We can solve the differential equations using Laplace transforms.

Let *f*(*x*) be a function defined for *x* > 0. The Laplace transform of *f*(*x*) is a function of *σ* as follows:

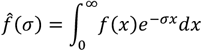

The Laplace transform of the derivative of a function *f*(*x*) is:

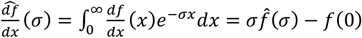

where a partial integral is adopted. This equation indicates that the Laplace transform of the function is multiplied by *σ* minus a constant. The constant is equal to the initial value of the original function.

Noting *L_k_*(0) = 0, we have the Laplace transform of the derivative of function *L_k_*(*x*) as:

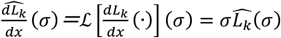

Note that the operation of calculating the derivative is the multiplication of *σ* to the Laplace transform of the original function. From this property, solving differential equations becomes the same as solving algebraic equations, the latter of which can be much simpler in its arithmetic.

The Laplace transform of Eq. (A.1a) becomes:

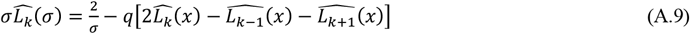

defined for *k* = 1,2,..,*n* – 1.

Using matrix *N* given in Eq. (A.2), Eq. (A.9) becomes:

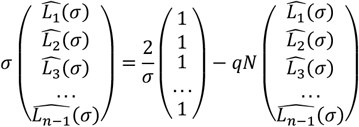

which leads to the following:

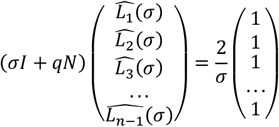

which is rewritten as:

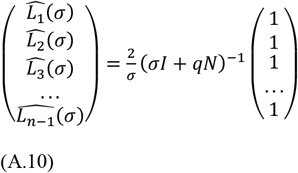

Hence once we obtain the inverse matrix in Eq. (A.10), we can obtain the original functions by an inverse-Laplace transform.

We illustrate this procedure for the case of *n* = 4. We set

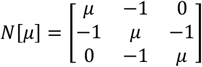

Its inverse matrix is:

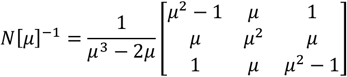

We can calculate as follows:

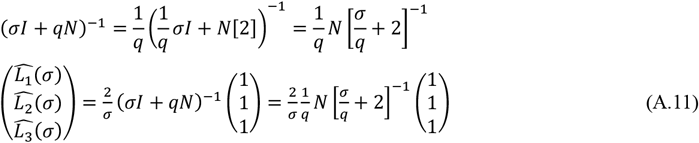

We note the following:

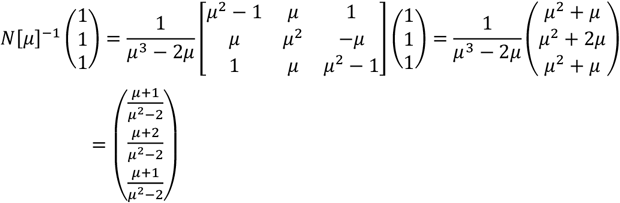

Because:

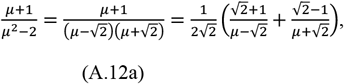

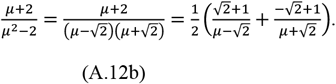

Using these, Eq. (A.10) is rewritten as:

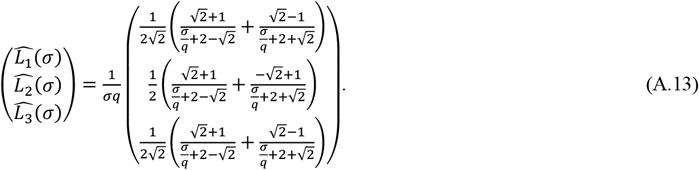

Here, we note 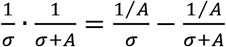 We have:

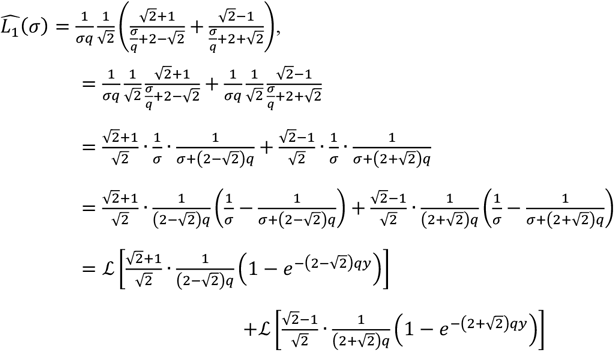

From this expression, we obtain:

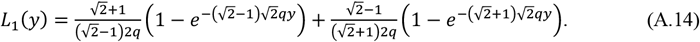

Note that *L*_3_(*x*) = *L*_1_(*x*), owing to the symmetry of the system with respect to *k*.

Similarly, we have:

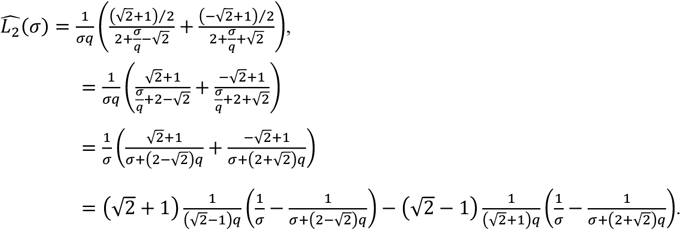

Inverse Laplace transform results in:

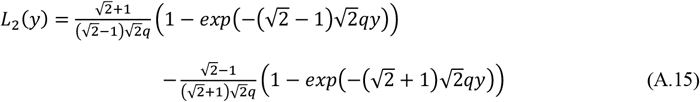

These results, given in Eqs. (A.14) and (A.15), are confirmed by the direct numerical analysis of the differential equations.

When *n* = 5 or larger, the same procedure is applicable. However, the inverse matrix becomes more complex, and the partial fraction expansion procedure becomes messy. We can at least assure that the inverse Laplace transform can generate an explicit solution of *L_k_*(*y*).

## Appendix B

### Solving a set of differential equations given in Eq. (3)

We consider the following square matrix of size *n* × *n*:

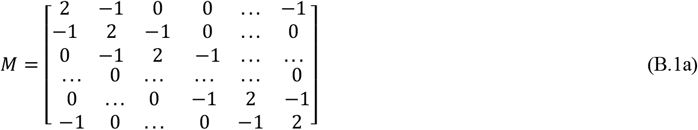

The elements of which are:

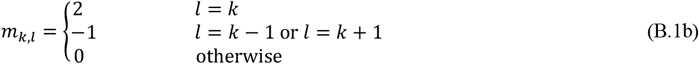

for *k* = 1,2,..,*n*. Because the circular boundary condition is adopted, *k* = 0 is the same as *k* = *n*.

Using matrix *M*, a system of differential equations given in Eq. (3) can be written as follows:

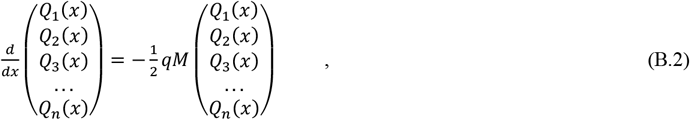

where *q* is the rate of stem cell replacement per cell per unit shoot length. If this event happens, the site occupied by the stem cell is filled by one of the nearest neighbors (either left or right, with equal probability). It is the probability of a stem cell exchange in a short time interval corresponding to the unit spatial length of the shoot. The initial distribution is *Q_n_*(0) = 1, and *Q_k_*(0) = 0 for *k* ≠ *n*. This is represented by a column vector in which the *n*th element (which is the same as the 0th element) is 1 and all the other elements are 0.

#### B.1. Solution using an exponential function of matrix M

We define an exponential function of a matrix as follows. Let *A* be a square matrix of size *n* × *n*. We consider an exponential function of *A* as follows:

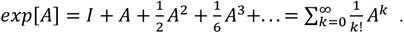

If we define:

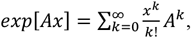

this quantity satisfies the following differential equation:

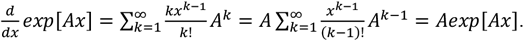

Hence, the solution of the differential equation (A.2) is written as:

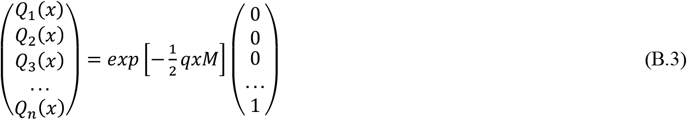

The rightmost vector is the initial distribution. We can confirm that Eq. (A.3) satisfies Eq. (A.2) and the initial condition.

#### B.2 Diagonalization of matrix M

To solve Eq. (B.3), we calculate an exponential function of matrix *M* in Eq. (B.3) in the following.

Let 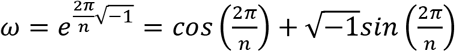. We have *w^n^* = 1. We consider the following *n* column vectors.

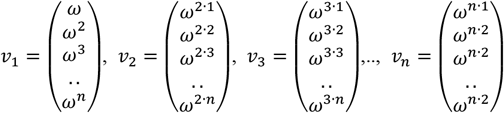

These vectors, *v*_1_, *v*_2_,.., *v_n_*, are eigenvectors of matrix *M*. We can show this by performing direct calculation:

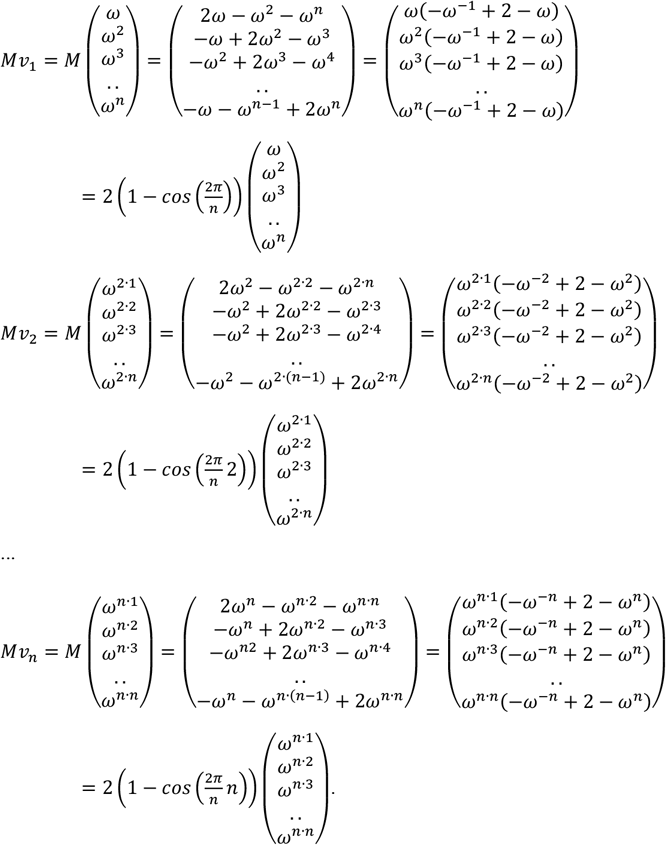

We note that 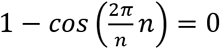. All the elements of the eigenvector corresponding to eigenvalue 0 are 1.

Hence, we have all the eigenvalues and corresponding (right-) eigenvectors. We consider a matrix in which all the columns are right eigenvectors for *M*.

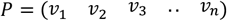

is a square matrix of size *n* × *n*. The (*k, l*)-element of matrix *P* is the *k*th element of vector *v_l_*. Hence,

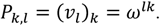

If we multiply matrices *M* and *P*, we have

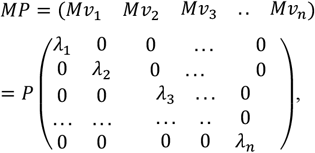

where 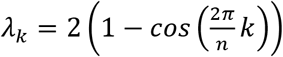. We multiply both sides by the inverse matrix of *P* from the right. The above equation becomes

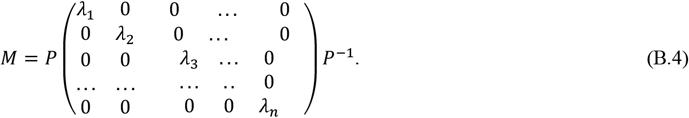

#### B.3. Exponential function of matrix M

If *f*(*x*) is a function of *x* that can be calculated using arithmetic and the converging sum of terms, we can define a function of matrix *M*. The results are as follows:

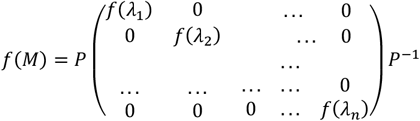

Next, we obtain the inverse of matrix *P*. We consider the transverse of matrix *P* and then change the complex conjugate for all the elements as follows:

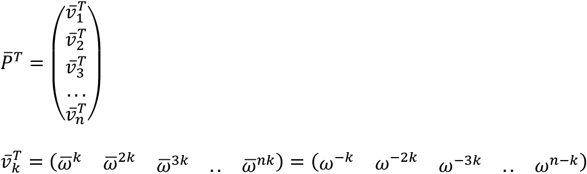

is a row vector and is the left eigenvector of matrix 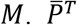 is a square matrix of size *n* × *n*. The corresponding eigenvalue is 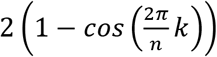. Here we note

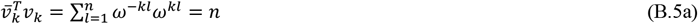

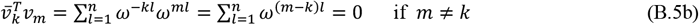

From these, we can obtain the inverse matrix as 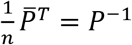.

The (*k, l*)-element of the inverse *P*^-1^ is:

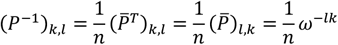

#### B.4. Solution of differential equations

Hence, we can rewrite the right-hand side of Eq. (A.3) as follows:

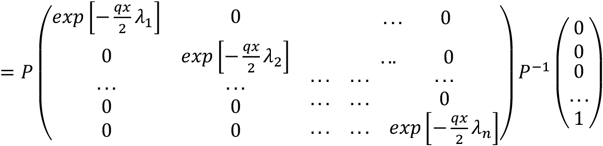

where 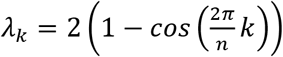 with *k* = 1,2,3,..,*n*. Note that 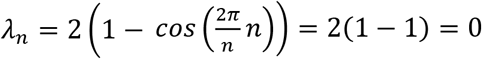. For 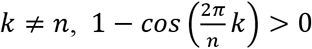 holds. Hence, 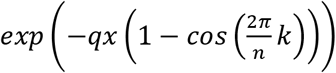 decreases with *x* and converges to 0 when *x* → ∞.

What remains should be the eigenvector corresponding to an eigenvalue of 0. This is the distribution with 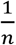 for all the elements. Hence, we have:

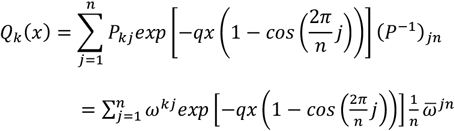

Because 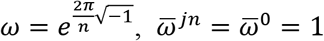 holds. We have:

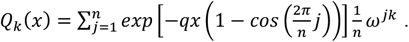

Here we note that 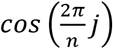 is the same if we replace *j* with −*j*. Note 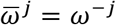.

The above equation is the same if we add the one with *j* replaced by −*j* divided by 2.

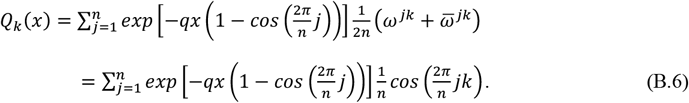

## Appendix C

### Perfect mixing case (Moran model)

In theoretical analyses of genetic diversity among stem cells in the meristem, starting from Klekowski and Kazarinova-Fukshansky (1984a), none of the research has addressed the possible effect of the angle around the shoot axis. Many models assume the Wright-Fisher process, with a synchronized replacement of the stem cells at the shoot apical meristem in a single population. The number of stem cells was fixed (often denoted by *α*). The stem cells in a generation *t* are produced from their parental generation *t* – 1 by random sampling.

Here, we calculate the corresponding Moran model with a continuous-time version in which the replacement of stem cells occurs asynchronously. At random times, one of the *n* stem cells is chosen and starts to divide. This results in one stem cell and one differentiated cell. The former remains in the stem cell population, but the latter is placed in one of the columns and becomes a cell of number “1”, with all the other cells in the same column becoming numbered shifted by one. This is in effect the same as the case in which *n* stem cells are not assigned to columns but they are in a mixed population at the center of the meristem.

Let *L*(*y*) be the mean coalescent length between two different cells at the same position along the shoot. Then, the following recursive formula is satisfied:

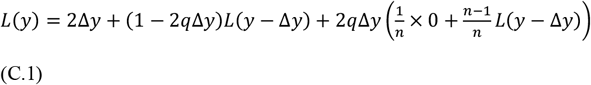

*c* is the probability for a stem cell to experience cell division in a unit length of time. If this is converted to the length of the shoot, *q*Δ*y* = (*c/rg*)Δ*y* is the probability for a single ancestral cell to experience cell division and replacement in a short period corresponding to shoot length Δ*y*. The first term on the right-hand side of Eq. (C.1) indicates the gain of coalescent length by a short length Δ*y*. It has a factor of 2 because there are two ancestral cells. The second term indicates the probability for no change to occur for the two sampled cells (1 – 2*q*Δ*y*) multiplied by the mean coalescent length between them at position *y* – Δ*y*. The third term on the right-hand side indicates the case with one of the two sampled cells experiencing cell division (with probability *2qΔy*). If this takes place, the newly born cell could be a daughter cell of the other sampled cell or not, with probability 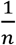 and 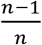, respectively.

Eq. (C.1) becomes

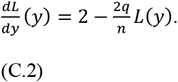

The initial condition is *L*(0) = 0 because all the stem cells are genetically identical at the beginning of the shoot elongation. Then Eq. (C.2) is solved as

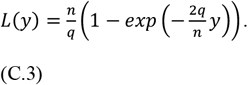

This is much smaller than the case in which stem cells at the shoot apical meristem have spatial structure. In the limit of *y* → ∞. It converges to 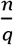 Because the coalescent length between the same cell is 0, the mean value of the coalescent length between two randomly sampled cells is 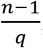, which is 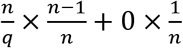. This argument is based on a continuous-time Moran process in which a stem cell is replaced by a randomly chosen cell among all the *n* stem cells including the one that failed to leave the successor (see Nowak 2006).

However, in the stem cell dynamics discussed in this paper, a stem cell is replaced by a copy of the other stem cells in the meristem. In choosing the cell that replaced it, we excluded the stem cell that failed to leave its successor stem cell. If we consider this situation we have a recursive formula that is similar to Eq. (C.1) but the brackets in the second term of the right-hand side should be replaced by 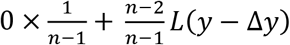. Then the differential equation is:

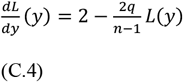

instead of Eq. (C.2). The solution is:

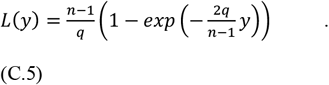

instead of Eq. (C.3). In the limit of *y* → ∞. It converges to 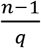. The mean value of the coalescent length is (*n* – 1)^2^/*nq*.

In contrast, the mean value of the coalescent length when replacement occurs only by the nearest neighbor in the limit *y* → ∞ is 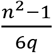. The latter is greater than the result of the Moran model 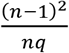 for *n* ≥ 4. The ratio is *n*(*n* + 1)/6(*n* – 1). In the Moran model, cell replacement occurs at every cell division. In contrast, in the model we discuss in this paper, stem cells normally engage in asymmetric division leaving their successor cells, and stem-cell replacement occurs only rarely after failure. In addition, the latter is much greater than the former when *n* is reasonably large. This effect can be understood as geographic differentiation in a stepping-stone model of 1 dimension (circular habitats) being larger than a completely mixed population with the same total population size (Maruyama 1970).

## Appendix D

We consider the situation in which stem cells occasionally undergo an exchange of locations between the nearest neighbors. Let s be the rate of stem-cell exchange per unit time per pair of nearest-neighbor stem cells. Let *q*’ = (*c* + *s*)/*rg*, and *q* = *c/rg*, satisfying 0 < *q* < f.

### D.1 Probability distribution of the ancestral stem cell for a stem cell

All the equations for *Q_k_*(*x*) remain the same as in the text if we replace *q* with *q*’ = (*c* + *s*)/*rg*. Appendix A also holds if we replace *q* with *q*’. The transition between states differing in *k* by one occurs faster, because not only stem-cell replacement but also stem-cell exchange, contributes to column shifting. The movement of the ancestral lineage in Fig. 7 is simply faster than before.

### D.2 Coalescent length between cells sampled at the same position

The equations for *L_k_*(*y*) are different for *k* = 1 (which is the same as *k* = *n* – 1) and for others (*k* = 2,3,…,*n* – 2). Eq. (2) becomes:

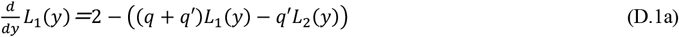

and

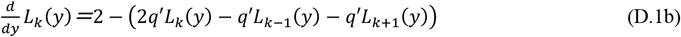

for *k* = 2,3,...,n – 2.

We note that both stem-cell exchange and stem-cell replacement cause a state change from *k* to *k* + 1 or to *k* – 1 if *k* = 1,2,…,*n* – 2. In contrast, the coalescent of two ancestral lineages occurs only by stem-cell replacement but not by stem-cell exchange. Hence, the transition from *k* = 1 to *k* = 0 occurs at rate *c*, which is lower than the transition for other state changes (such as from *k* = 1 to *k* = 2 or from *k* = 2 to *k* = 1) that occur at rate *c* + *s*.

In the limit *y* → ∞, we have

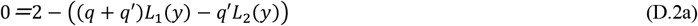

for *k* = 2,3,…,*n* – 2. The following solution:

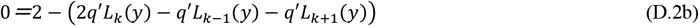

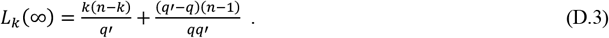

satisfies Eqs. (D.2a) and (D.2b). We can confirm this by direct replacement.

Eq. (D.3) is the same as Eq. (11) in the main text. Eq. (D.3) becomes Eq. (3) when *s* = 0, which makes *q*’ = *q*.

When stem cells are very mobile and the exchange rate is fast (*s* is large), *q*’ → ∞ holds, and Eq. (D.3) becomes 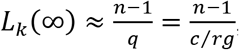, which is independent of angle *k*. The genetic distance between the different stem cells becomes independent of the angle. It can be large if the stem-cell replacement rate *c* is small. See Fig. 8B for an illustration.

If we compare the current model where replacement takes place only between nearest neighbors, and the Moran model in which replacement can be done by any other stem cells (studied in Appendix C), the result in the limit *q*’ → ∞ is the same. Hence, the random exchange of neighboring stem cells takes place very fast, and the model is basically the same as the Moran model if every stem cell division is accompanied by a stem cell replacement.

